# Light spectra trigger divergent gene expression in barley cultivars

**DOI:** 10.1101/2021.02.03.429565

**Authors:** Arantxa Monteagudo, Álvaro Rodríguez del Río, Bruno Contreras-Moreira, Tibor Kiss, Marianna Mayer, Ildikó Karsai, Ernesto Igartua, Ana M. Casas

**Affiliations:** Aula Dei Experimental Station (EEAD-CSIC), Avda. Montañana 1005, E-50059 Zaragoza, Spain; Fundación ARAID, Zaragoza, Spain; Centre for Agriculture Research ELKH (ATK), H-2462 Martonvásár, Hungary

**Keywords:** Barley, cold-regulated genes, development, light quality, RNA-seq, senescence, signaling, transcription factors

## Abstract

Light spectra influence barley development, causing a diverse range of responses among cultivars that are poorly understood. Here, we exposed three barley genotypes with different light sensitivities to two light sources: fluorescent bulbs, over-representing green and red wavebands, and metal halide lamps, with a more balanced spectrum. We used RNA sequencing to identify the main genes and pathways involved in the different responses, and RT-qPCR to validate the expression values. Different grades of sensitivity to light spectra were associated with transcriptional reprogramming, plastid signals, and photosynthesis. The genotypes were especially divergent in the expression of genes regulated by transcription factors from MADS-box, WRKY, and NAC families, and in specific photoreceptors such as phytochromes and cryptochromes. Variations in light spectra also affected the expression of circadian clock, flowering time, and frost tolerance genes, among others, resembling plant responses to temperature. The relation between *PPD-H1, HvVRN1*, and *HvFT1* expression might explain genotypic differences. Light-sensitive genotypes experienced a partial reversion of the vernalization process and senescence-related stress under the less favorable light quality conditions. The observed light-quality sensitivities reveal a complex mechanism of adaptation to regions with specific light quality features and/or possible regulation of light spectra in plant development during early spring.

**Highlight:** Development genes were affected by light quality in the barley varieties tested. Different grades of sensitivity were related to the expression of transcription factors, senescence, light signaling and cold-regulated genes.

## Introduction

As sessile organisms, plants have evolved adapting and surviving in a wide variety of environments. One of the main developmental triggers is light, whose features regulate growth and determine the adaptation to changing environments with different light duration, quantity, and quality (Franklin, 2009; Ugarte *et al*., 2010).

Like day length, light quality and intensity are not constant in natural environments, as revealed by the different spectra that occur in different moments of the day, seasons, climates, and atmospheric conditions (Holmes and Smith, 1977; Smith, 1982). Although the responses of plants to some light features have been thoroughly analyzed (Franklin, 2009; Ugarte *et al*., 2010; Monostori *et al*., 2018), there is a gap in the study of natural genetic variation in crop plants and its possible effect on crop development and adaptation. For instance, light quality effects have been widely studied in *Arabidopsis* (Adams *et al*., 2009), but only to a lesser extent in cereals (Ugarte *et al*., 2010).

Differential regulation of genes whose expression levels are affected by light quality signals may reflect different abilities to compete in diverse species or crop varieties. The concentration and efficiency of photoreceptors that control the dynamics of red/far-red (R/FR) light signaling are highly variable in different species and crop varieties (Merotto Jr. *et al*., 2009). Higher plants possess two types of signal-transducing photoreceptors: phytochromes (in cereals PhyA, PhyB, and PhyC) absorbing principally in the 600-800 nm waveband, and cryptochromes (Cry1, Cry2), absorbing only in the 300-500 nm band (Smith 1982; Casal 1993). Phytochrome proteins are characterized by a red/far-red photochromicity, changing their spectral absorbance properties upon light absorption. Their biologically inactive form activates after absorbing R light and reverts to inactive after absorbing FR light (Rockwell *et al*., 2006). As dimeric proteins, the role of homo and heterodimers is still an open area of research but it has been proven that both *PhyB* and *PhyC* genes are required for the induction of wheat flowering under long photoperiods (Pearce *et al*., 2016). Cryptochromes have been found to regulate photomorphogenesis and the expression of genes involved in blue light signaling and stress response (Kleine *et al*. 2007). Both phytochrome dimers and cryptochromes interact with transcription factors (TF) known as Phytochrome Interacting Factors (PIFs) (Leivar and Monte, 2014; Pedmale *et al*., 2016), regulating clock and flowering time genes (Oakenfull & Davis 2017), and are at the top of fundamental light-driven processes.

Together with light, crops responses to temperature are receiving increasing attention, and more efforts are being dedicated to unraveling the catalog of cross-talk and nodes at which both signals converge (Franklin *et al*., 2014). Both are particularly important in winter cereals, which need to satisfy cold needs (called *vernalization*) before the spring when long days (LD) mark the signal to flower. Thus, perception of photoperiod and cold are critical to enabling flowering timely. Two main genes control the vernalization response in winter cereals. In barley, these genes are *HvVRN1*, an *AP1*-like MADS-box TF, and promoter of flowering, and *HvVRN2*, also known as *ZCCT-Ha-c*, a zinc-finger and CCT domain-containing repressor protein that belongs to the *CONSTANS*-like family (Trevaskis *et al*., 2003; Yan *et al*., 2004). Both interact with the floral pathway integrator *HvFT1* (Yan *et al*., 2006). When the cold requirement has not been satisfied, long-days promote *HvVRN2* expression, repressing *HvFT1*, and delaying flowering until plants complete vernalization (Trevaskis *et al*., 2006; Hemming *et al*., 2008). In temperate cereals, other members of the MADS-box TF family genes, such as the *Flowering Locus C* (FLC)-clade member *OS2* (*ODDSOC2*) and the *Short Vegetative Phase* (*SVP*)-clade member, *VRT2*, cause a delay until experiencing enough cold (Kane *et al*., 2005; Greenup *et al*., 2010; Xie *et al*., 2019). Cold induces *HvVRN1*, which then represses *HvVRN2*, and together with the influence of LD, allows the expression of the flowering integrator *HvFT1*. During the vernalization of winter genotypes, the expression of the *SVP*-clade MADS-box TFs (*HvVRT2, HvBM1*, or *HvBM10*) is up-regulated during the vegetative phase (Trevaskis *et al*., 2007). Particularly, VRT2 participates in regulating the vernalization flowering pathway, interacting and cooperating with VRN1 (Xie *et al*., 2019), and after the transition to the reproductive phase, its expression declines (Kane *et al*., 2005). The connection between photoreceptor and photoperiod pathways has been attributed to PhyC, which activates the long-day photoperiod response gene, *PPD-1* in LD (Nishida *et al*., 2013; Chen *et al*., 2014; Pankin *et al*., 2014; Woods *et al*., 2014). Both *PhyC* and *PhyB* promote flowering under LD, although *PhyB* regulates more genes than *PhyC* (Kippes *et al*., 2020), particularly those involved in vegetative development, hormone biosynthesis, and signaling, shade avoidance response, abiotic stress tolerance (Pearce *et al*., 2016), and cold tolerance (Franklin and Quail, 2010; Novák *et al*., 2016).

Motivated by the lack of studies on natural genetic variation in crop plants and its possible effect on crop development and adaptation, we explored the phenotypic variability for plant growth, in response to different light quality environments, in barley (*Hordeum vulgare*, L.). Plants were exposed to two light sources: fluorescent light, which presents high peaks at the 550-650 nm regions, corresponding to green and red wavebands, and metal halide bulbs, which yield a more balanced spectrum. Although all genotypes had been vernalized before the start of the experiment and temperature, photoperiod and light intensity were identical and inductive to promote flowering, plants grown under fluorescent light showed delayed development compared to those under metal halide light (Monteagudo *et al*. 2020). We thus observed an effect of light quality on the expression of flowering time genes, opening new questions about the regulation of photoperiod and vernalization pathways in different barley varieties.

Here, we exposed three of the previously assessed genotypes to the same two contrasting light spectral conditions to further investigate their gene expression patterns: an insensitive line (minor response to light quality), Esterel, and two sensitive lines (major growth differences between light quality conditions), Price (sensitive-intermediate) and WA1614-95 (sensitive-extreme). The aims of this study were: a) to identify genes that explain the different sensitivities of barley lines to light spectral quality, and b) to gain further knowledge on the influence of light quality on development and on the expression of flowering time genes. Here we provide evidence of the molecular mechanisms affected by light quality. We identified key flowering pathway regulators, and other important genes involved in development whose expression was affected in all varieties. Genotypic differences in the expression of light signaling and cold responses were also found, which might be related to barley adaptability to a wide range of environments and to an additional regulation mechanism of plant development during early spring.

## Materials and Methods

### Plant material and phenotyping

We selected varieties Esterel, Price, and WA1614-95 (Supplementary Table S1), from a set of 11 winter or facultative barley varieties described in a previous study (Monteagudo *et al*., 2020) that revealed different responses to light environments (see Figure 1).

**Figure 1.**
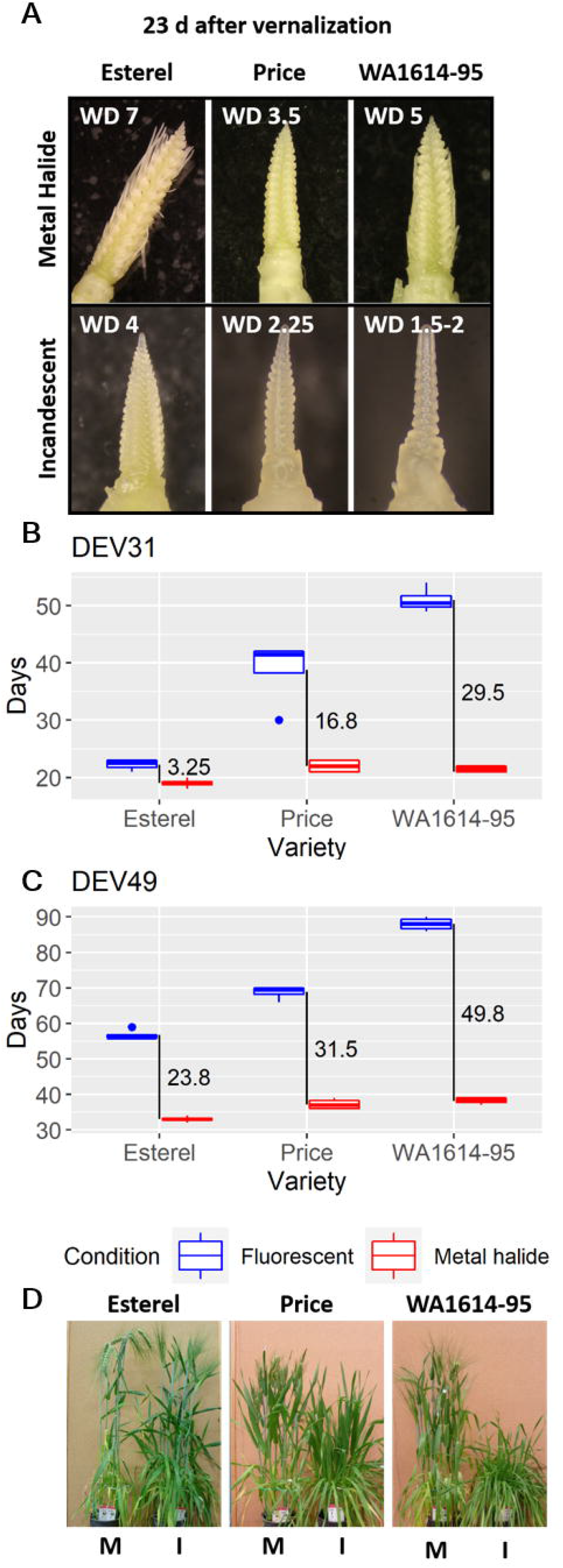
Phenotypic differences between varieties. A) Apex development in plants dissected 23 days after the end of the vernalization treatment. WD, Waddington stage. B) and C) Days to first node (DEV31) and awn appearance (DEV49), expressed in days from the end of the vernalization treatment, measured in 4 biological replicates. Vertical black lines represent the days of difference between fluorescent and metal halide light conditions. D) Plants photographed 58 days after the end of the vernalization treatment.

The experiment was carried out at the phytotron facilities of the Agricultural Research Institute of the Hungarian Academy of Sciences, Martonvásár (Hungary), using Conviron PGR-15 growth chambers (Conviron Ltd., Canada). Two light treatments were established in independent growth chambers. Two lamp types were used: Sylvania cool white fluorescent (F) and Tungsram HGL-400 metal halide (M) light bulbs. The height of the lamps in the F chamber was adjusted once per week to 1.4 m above the canopy, to match the light intensity of the M chamber, in which the lights were set at a fixed height. The conditions in both chambers were set to long photoperiod (16 h light/8h night), and 18 ± 1°C constant temperature, and light intensity of 250 µmol m^-2^ s^-1^. Temperature was continuously monitored through an air-sampling channel, located in the middle of the cabinet at canopy level. This system of temperature control eliminated the possibility that plants experienced different temperatures at both chambers.

For phenotypic measurements, four seeds per genotype and treatment were sown in individual pots (12 × 18 cm, 1.5 kg). Additionally, 20 seeds per genotype and treatment, sown in groups of 5 plants/pot, were used for destructive samplings to record apex development stage, and for gene expression studies. All plants were fully vernalized (5 ± 2°C for 52 days under 8h light/16 h night, low-intensity metal-halide light bulbs) before entering the light quality chambers, to synchronize the development of the three genotypes. Plant development was monitored twice a week, checking for first node appearance (plant developmental stage 31, or DEV31) and appearance of the awns just visible above the last leaf sheath (DEV49). All these data were defined based on stages of the Zadoks’s scale (Zadoks *et al*., 1974), following the description of Tottman *et al*. (1979). Apex dissection was carried out 23 days after the end of the vernalization period in 3 plants per variety and treatment. Phenotyping consisted of recording apex stage following the Waddington’s scale (Waddington *et al*., 1983). Plants were grown to full maturity.

### RNA extraction and transcriptome sequencing

Three genotypes were used for transcriptome analysis. Three biological replicates per genotype and treatment were produced. Each biological replicate pooled the last expanded leaves of the main tillers from two different plants. Leaves were sampled in the middle of the light cycle, 20 days after the end of the vernalization period, and immediately frozen in liquid N_2_. Leaf samples for Real-time PCR quantification (qRT-PCR) validation were obtained in an independent experiment, as reported by Monteagudo *et al*. (2020).

Total RNA was isolated with TRIzol (Thermo Fisher Scientific, Ltd.) followed by the Qiagen RNeasy plant mini kit, following the manufacturer’s instructions (Qiagen, Ltd.). Then, the material was extracted in the QIAcube equipment (Qiagen, Ltd.) with an extra step of DNase treatment programmed. RNA quality was assessed with a NanoDrop 2000 spectrophotometer (Thermo Fisher Scientific, Ltd.) at ATK (Hungary).

RNAseq was performed by Novogene (HK) Co. Ltd. (China) after quality controls by agarose gel electrophoresis and Bioanalyzer 2100 (Agilent, USA; RIN ≥ 6.3). Library construction was developed from enriched RNA, using oligo(dT) beads. Then, mRNA was randomly fragmented, followed by cDNA synthesis using random hexamers and reverse transcriptase. After the first-strand synthesis, a custom second-strand synthesis buffer (Illumina, USA) was added, with dNTPs, RNase H, and *Escherichia coli* polymerase F to generate the second strand by nick-translation, and AMPure XP beads were used to purify the cDNA. The final cDNA library was ready after a round of purification, terminal repair, A-tailing, ligation of sequencing adapters, size selection, and PCR enrichment. Library concentration was first quantified using a Qubit 2.0 fluorometer (Life Technologies) and then diluted to 1 ng/μl before checking insert size on an Agilent 2100 and quantifying to greater accuracy by quantitative PCR (library activity > 2 nM). Eighteen barcoded libraries were multiplexed and sequenced, 2×150 bp paired-end reads, in an Illumina HiSeq^™^ 2500 sequencer, yielding on average 50 Million reads per sample. The whole dataset consisted of 18 samples, i.e. 3 biological replicates, from 3 varieties and 2 light conditions.

Raw reads were processed with Illumina CASAVA v1.8 (Illumina, USA). Low-quality reads (reads with more than 50% low-quality base (Q ≤ 20)) were removed. Reads from the three genotypes were jointly assembled *de novo*, with the software Trinity (Haas *et al*., 2013) (Figure 2). The obtained assembled transcripts had a median length of 366 bp. Raw transcripts of all three genotypes were combined, followed by a step of hierarchical clustering. Then, the longest transcripts were kept and unigenes were called with Corset v1.05 (-m 10 to remove redundancy, Davidson and Oshlack, 2014). The median unigene length was 779 bp. Raw reads of the sequencing experiments (accessions ERR3763262-ERR3763279) and assembled *de novo* transcripts (ERZ1264422) have been submitted to the European Nucleotide Archive.

**Figure 2.**
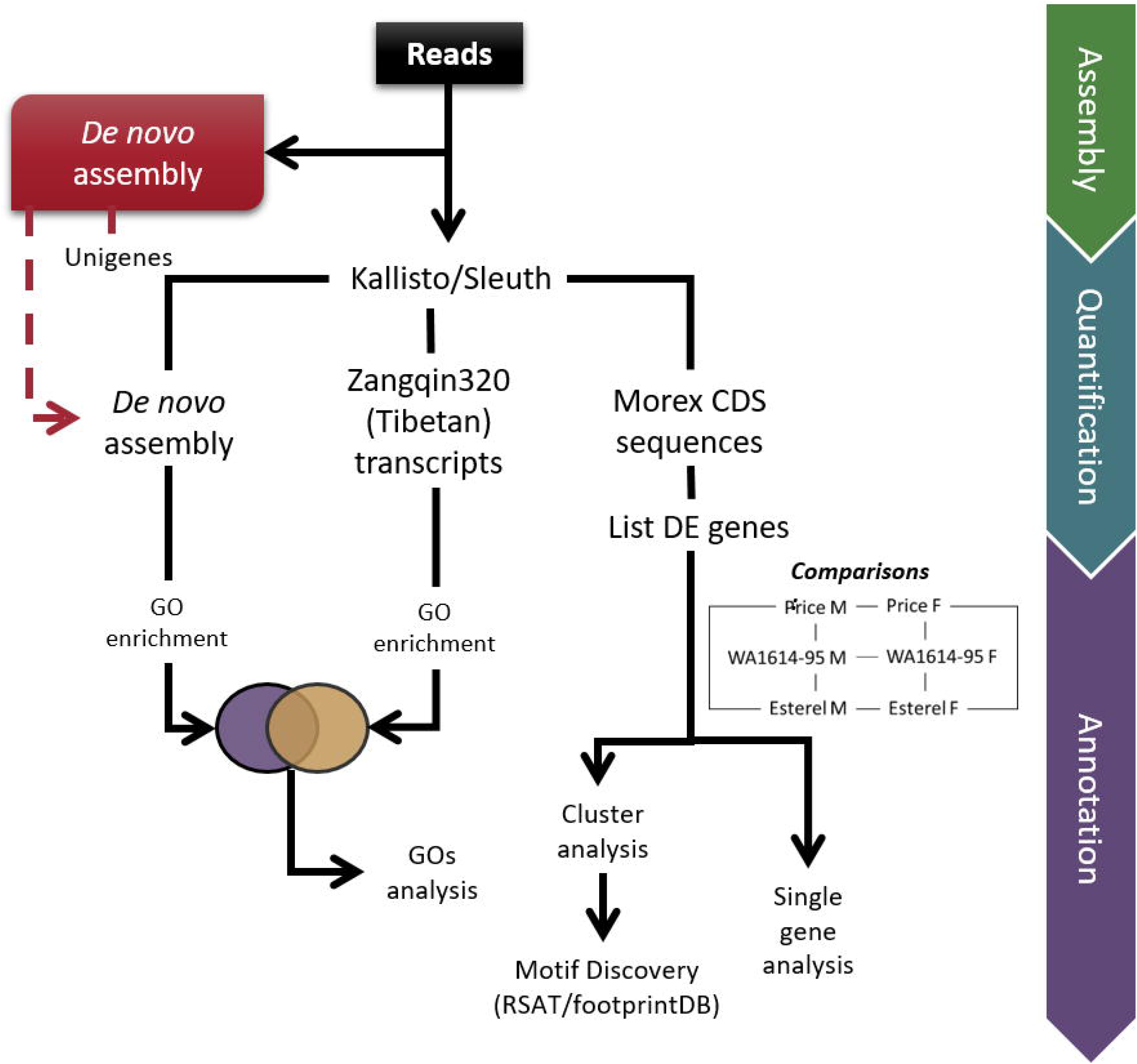
Pipeline of the RNA-seq analysis. Because of the incomplete GO annotation of the Morex 2017 genome (several GO terms involved in light stress are missing), we also calculated DE genes using the two best references according to our benchmarking, for performing the GO enrichment analysis.

### Quantification of gene expression and differential expression analysis

There are several barley assembly references available. We tested eight references and chose the most suitable for each purpose (Supplementary Information). We mapped clean reads against each reference and quantified transcript abundance as Transcripts Per Million (TPM) using Kallisto (Bray *et al*., 2016). We used the R functions ‘heatmap’ and ‘hclust’ (R Core Team, 2020), to cluster the gene expression patterns from the experimental replicates, from the three genotypes, under the two light conditions. The resulting clusters were used to assess the expression quantification with several mapping references (see Supplementary Information). The gene sets which grouped more biological replicates together were selected for gene expression quantification. The barley reference genome from cultivar Morex (named **“Morex CDS sequences”**) (Mascher *et al*., 2017) was used for all downstream analyses except GO enrichment, as these sequences lacked relevant GO terms related to light responses. For this reason, assembled Zangqing320 PacBio reads, from here on noted as **“Tibetan transcripts”** (Dai *et al*., 2018), and the assembly generated in this work (**“*de novo* assembly”**), were chosen for conducting the GO enrichment analysis (Figure 2).

For ensuring the quality of the biological replicates, we calculated Pearson correlation coefficients across the transcript abundance of biological samples and replicates derived from reads mapped to **“Morex CDS sequences”**, with the R package “corrplot” (Wei and Simko, 2017). Independently validated RNA-seq expression values were obtained through qRT-PCR (ABI 7500, Applied Biosystems), performed on biological replicates for the same varieties and conditions, grown in an independent experiment. Expression values of 10 genes (Supplementary Table S2) were calculated relative to *Actin* (*HORVU5Hr1G039850*.*3*), taking into account the efficiency of each pair of primers. Each PCR reaction contained 5 µl of PowerUp SYBR Green Master Mix (Applied Biosystems), 0.5 µM of each primer, and 250 ng of cDNA in a volume of 10 µl. Reactions were run with the following conditions: 2 min activation at 50°C, 10 min of pre-denaturation at 95°C; followed by 44 cycles of 15 s denaturation at 95°C, 50 s annealing at 60°C, and 45 s of extension step at 72°C, ending with a melting curve 60°–95°C default ramp rate. The same normalization (relative to *Actin*) was accomplished for the number of TPM values and compared for the same genes and treatments in the RNA-seq. This procedure was carried out with a total of 60 points (average of 3 biological replicates per treatment and variety, in 10 genes).

We used Sleuth (Pimentel *et al*., 2017) for calculating differential expression (DE) of the genes from the three barley varieties. Nine DE analyses (M vs F for each one of three genotypes, and comparisons between pairs of genotypes for each light quality condition) were achieved for each mapping reference. However, here we only report results on the DE genes calculated from the three aforementioned references (Figure 2 and Supplementary Information). We validated the Kallisto/Sleuth methodology by comparing it to the RSEM (Li and Dewey, 2011) and DESEQ (Anders and Huber, 2010) pipelines, originally used by Novogene (Supplementary Information).

DE isoforms were detected using False Discovery Rate (FDR) adjusted p-values (named hereby “q-values”), setting the threshold at 0.05. As plants responded better to metal halide light, such condition was considered as control. Thus, DE genes are expressed in terms of being up- or down-regulated under fluorescent light.

We used the DE genes calculated with Morex CDS reference for describing the patterns of expression of individual genes. To reduce the number of DE genes to a workable number and increase confidence, we only focused on DE genes with q-value < 0.01 (“Key genes”) for producing Venn diagrams and the identification of individual genes affected by light quality. All the key genes were referenced to the barley reference genomes Morex v1 (Mascher *et al*., 2017), and Morex v2 (Monat *et al*., 2019).

### GO enrichment analysis

We used DE genes (q-value < 0.05) calculated from the “Tibetan transcripts” and “*de novo* assembly” references in three within genotype comparisons for the GO enrichment tests. We mapped the three sets of DE genes calculated from both references against the Morex genome assembly WGS (Mayer *et al*., 2012) in PlantRegMap (Jin *et al*., 2017), which uses reciprocal best BLAST hits to assign Morex gene *id*s to query sequences. Matched genes received the GO terms from the Morex reference. Enrichment analysis was calculated by Fisher’s exact test, with the complete gene set of Morex as control. GO terms with a q-value < 0.05 were considered enriched. GO term enrichment analysis was performed independently for the three sets of DE genes derived from the two mapping references; we considered GO terms enriched in both references for each of the three within genotype comparisons as highly reliable.

### Clusters of DE genes

From within genotype comparisons in the two light conditions, we selected DE genes obtained using the Morex CDS sequences as mapping reference. TPMs values were clustered by the k-means method using the ‘eclust’ function from the ‘factoextra’ R package (Kassambara and Mundt, 2020), which separates the points into a defined k number of groups and returns the total within-cluster sum of squares. The optimal number of clusters was calculated minimizing and stabilizing that term.

### Motif discovery

We manually selected some clusters based on their different expression patterns between light conditions in sensitive and insensitive varieties. To retrieve promoter sequences of the corresponding clustered DE genes, we extracted upstream sequences (−500, +200 nucleotides around annotated Transcription Start Sites) for each gene, from the server http://plants.rsat.eu (Nguyen *et al*., 2018) and performed the motif discovery protocol described in Contreras-Moreira *et al*. (2016) and Ksouri *et al*. (2021). For each cluster analyzed, 50 clusters of the same size made by random picking upstream barley sequences were used as negative controls for assessing the significance of motifs found (parameters MAXSIGGO=60 MAXSIG=10 MINCOR=0.7 MINNCOR=0.5). The resulting motifs were compared to motifs annotated in the footprintDB database (Sebastian and Contreras-Moreira, 2014). The complete motif discovery results are available at http://rsat.eead.csic.es/plants/data/light_report.

## Results

### Diversity in the response to different light sources

The three varieties differ in growth habit. The *HvVrn2* gene is present in Esterel and Price and absent in WA1614-95 (Supplementary Table S1), and all three genotypes have a winter allele at *HvVrn1*. Therefore, Esterel and Price are winter varieties, whereas WA1614-95 is a facultative variety. The vernalization treatment placed the three lines at a similar developmental stage (between Z11 and Z12) at the beginning of the light treatments. Under M conditions, development was accelerated, compared to F, as revealed by the more developed apices in 23-day old plants (Figure 1A). All three varieties flowered earlier in M than in F. However, Esterel showed the least differences between treatments in days to the appearance of the first node (DEV31, Figure 1B) and days to awns appearance (DEV49, Figure 1C, and D). WA1614-95 presented the largest differences, with Price in an intermediate position (Monteagudo *et al*., 2020).

### RNA-seq performance

Sampling for RNA-seq took place three days before the examination of the apices shown in Figure 1. At that time, all three varieties had started the reproductive phase, or at least were very close to reaching that point. Sequencing cDNA of 18 samples produced a total amount of 1.92 billion paired-end reads. The joint *de novo* assembly for the three genotypes contained 375,488 isoforms, from which we obtained 181,337 unigenes. We benchmarked different barley references for mapping reads to transcripts (Supplementary Information). As Morex is the most widely used barley reference, performs well in our reference benchmarking, and has been functionality annotated by the community over the years, “Morex CDS sequences” were chosen to calculate gene expression values and calling DE genes (Figure 2). However, because of the incomplete GO annotation of the Morex 2017 genome (for instance, several GO terms involved in light stress were missing), we also calculated DE genes using two of the best references according to our benchmarks (Supplementary Information): our assembly (“*de novo* assembly” DE genes) and the “Tibetan transcripts” (Dai *et al*., 2018). Note that DE genes calculated from these two references were only used for GO enrichment analysis.

We used correlation coefficients between genes, estimated counts across the three biological replicates, as a sample quality control. Esterel_F2 showed lower correlation coefficients with the other two biological replicates than the remaining samples (Supplementary Figure S1B). Consequently, replicate Esterel_F2 was discarded from downstream analyses. Additionally, it is remarkable that Price and WA1614-95 expression patterns were highly correlated in M conditions, consistently for all replicates (Supplementary Figure S1A), but showed lower correlation coefficients in F conditions (Supplementary Figure S1B), indicating similar responses of these two genotypes to M light, but variable responses when exposed to fluorescent light. Esterel (insensitive genotype) showed lower expression level correlation coefficients with the other genotypes in both conditions.

### Expression analysis of key genes

To unravel differential responses of the three genotypes to light quality, we focused on DE genes occurring within each genotype across light treatments. Price, WA1614-95, and Esterel initially showed 2,869, 4,218, and 3,591 DE genes with q-value < 0.05, listed in Supplementary Datasets S3, S4, and S5. To focus on genes most likely affected by light conditions, we further filtered the list considering only genes with q-value < 0.01 for subsequent analysis, and denoted them as “key genes”. Key DE genes in Esterel were predominantly down-regulated (in F compared to M), whereas Price showed more up-regulated than down-regulated genes, and WA1614-95 showed a similar number of up- and down-regulated DE genes (Figure 3).

**Figure 3.**
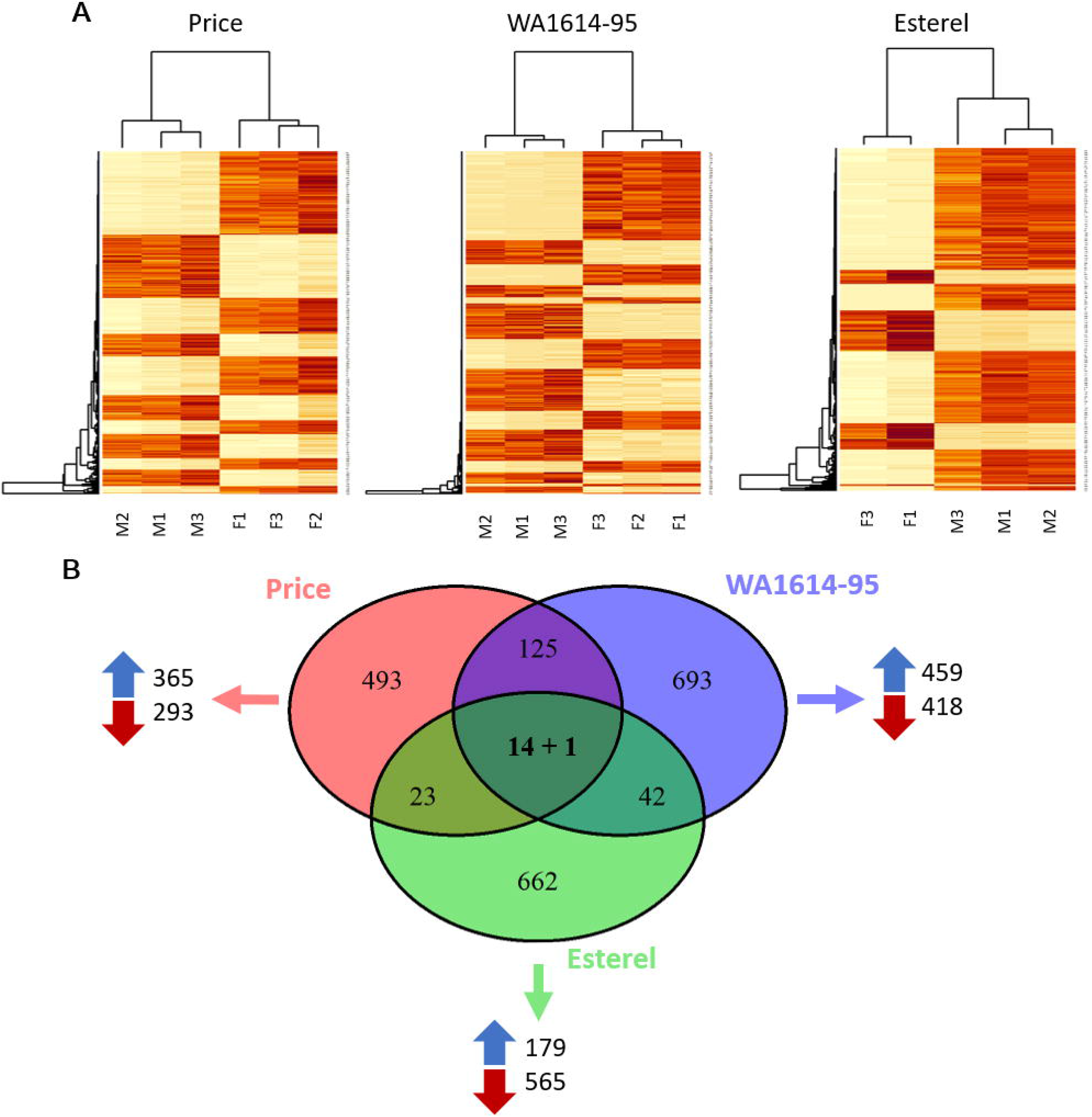
Differentially expressed (DE) genes in the three genotypes (q-value < 0.01). A) Clustering of DE genes with Morex CDS sequences used as reference. Colour code: red, upregulated in fluorescent light; yellow, downregulated. Three biological replicates per variety and condition are represented, except for Esterel in fluorescent light, from which a replicate was discarded. F, fluorescent; M, metal halide. B) Venn diagram showing the intersection in the number of DE genes among genotypes. Blue and red arrows indicate tthe number of DE genes upregulated or down regulated in fluorescent conditions in each genotype. The intersection between the three varieties indicates the number of DE genes that are annotated as High Confidence (14) and Low Confidence (1) genes in the Morex genome v2 (Monat *et al*. 2019).

The intersection of key genes for the three genotypes comprised 17 sequences (Table 1). Among them, *HvBM3* (*Barley MADS-box 3*), *HvBM8, PPD-H1* (*PSEUDO RESPONSE REGULATOR 7, HvPRR37*, Turner *et al*., 2005), *HvFT1* (*FLOWERING LOCUS T*-like, Yan *et al*., 2006) were down-regulated under fluorescent light conditions in the three genotypes, whereas *HvVRT2* (*VEGETATIVE TO REPRODUCTIVE TRANSITION 2*) and *RVE7*-like (*EARLY PHYTOCHROME RESPONSIVE 1*/*REVEILLE7*) were up-regulated in fluorescent light in the three genotypes. *RVE7*-like and *HvVRT2* were expressed at higher levels in WA1614-95, whereas *HvBM3, HvBM8*, and *HvFT1* showed higher expression in the insensitive line, Esterel (Figure 4). Two genes were annotated as *HvFT1*, aligned to chromosomes 3H and 7H, with the same expression levels. As *HvFT1* is located exclusively on 7H (Yan *et al*., 2006), we believe the hit on 3H probably comes from a duplication/misassembly in the Morex genome assembly v1, which contains a very large number of fragmented genes (Beier *et al*., 2017; Prade *et al*., 2018). After referencing key genes to the reference Morex v2 (Monat *et al*., 2019), we found that *HvBM8* was also duplicated (*HORVU2Hr1G063800* and *HORVU2Hr1G063810*, Supplementary Dataset S1), thus leaving 15 single copy key DE genes in the three lines (Figure 3). The 15 key genes were grouped in 14 high confidence (HC) and 1 low confidence (LC) genes according to their annotation. Only the LC gene (*Ethylene-responsive transcription factor*) followed different expression patterns among genotypes (up-regulated in F in the sensitive genotypes, Price and WA1614-95 and down-regulated in the insensitive genotype Esterel). This gene lacks a functional annotation in Morex v1 (only 159 bp were found in common between the 2017 reference and the RNA-seq sequences). Instead, in Morex v2 the gene has a larger coverage (539 bp). The remaining DE genes showed the same expression directions in all three genotypes.

**Table 1.** List of DE genes (q-value < 0.01) shared by three barley varieties studied. We show up-regulated (u) and down-regulated (d) genes in fluorescent light for each genotype.

| Target ID Morex v1.0<sup>a</sup> | Target ID Morex v2.0<sup>b</sup> | Description<sup>b</sup> | Gene | Cite | Price | WA1614-95 | Estrel |
| --- | --- | --- | --- | --- | --- | --- | --- |
| HORVU0Hr1G003020.3 | HORVU.MOREX.r2.2H G0105150.1 | MADS box transcription factor | HvBM3 | 1, 2 | d | d | d |
| HORVU2Hr1G063800.7 | HORVU.MOREX.r2.2H G0129220.1 | MADS box transcription factor | HvBM8 | 1, 2 | d | d | d |
| HORVU2Hr1G063810.1 |  |  |  |  | d | d | d |
| HORVU7Hr1G036130.1 | HORVU.MOREX.r2.7H G0551090.1 | MADS box transcription factor | HvVRT2 | 3, 4 | u | u | u |
| HORVU7Hr1G083670.3 | HORVU.MOREX.r2.7H G0591700.1 | Cytochrome P450 family protein |  |  | u | u | u |
| HORVU2Hr1G013400.32 | HORVU.MOREX.r2.2H G0088300.1 | Pseudo-response regulator | HvPRR37 (PPD-H1) | 5 | d | d | d |
| HORVU4Hr1G090860.12 | HORVU.MOREX.r2.4H G0348610.1 | Metacaspase-1 | cell death |  | u | u | u |
| HORVU5Hr1G071940.2 | HORVU.MOREX.r2.5H G0405790.1 | Glycosyltransferase |  |  | u | u | u |
| HORVU2Hr1G024120.10 | HORVU.MOREX.r2.2H G0097130.1 | Terpene synthase |  |  | d | d | d |
| HORVU0Hr1G038850.2 | HORVU.MOREX.r2.6H G0516180.1 | Protein kinase |  |  | u | u | u |
| HORVU3Hr1G111550.2 | HORVU.MOREX.r2.3H G0270770.1 | Ethylene-responsive transcription factor |  |  | u | u | d |
| HORVU3Hr1G021880.1 | HORVU.MOREX.r2.3H G0198440.1 | Glycosyltransferase | Glycosyl-transferase |  | d | d | d |
| HORVU3Hr1G087100.1 | HORVU.MOREX.r2.7H G0542540.1 | Flowering locus T | HvFT1 (VRN-H3) | 6, 7 | d | d | d |
| HORVU7Hr1G024610.1 |  |  |  |  | d | d | d |
| HORVU5Hr1G029260.1 | HORVU.MOREX.r2.5H G0372130.1 | Protein kinase family protein |  |  | d | d | d |
| HORVU2Hr1G104580.2 | HORVU.MOREX.r2.2H G0162680.1 | Homeodomain-like superfamily protein | RVE7-like |  | u | u | u |
| HORVU1Hr1G076460.3 | HORVU.MOREX.r2.1H G0062670.1 | Hemoglobin |  | 8 | u | u | u |

<sup>a</sup> Morex reference genome v1.0 (Mascher *et al.*, 2017)
<sup>b</sup> Morex reference genome v2.0 (Monat *et al.*, 2019)
1, Schmitz *et al.* (2000); 2, Ejaz and von Korff (2017); 3, Kane *et al.* (2005); 4, Szucs *et al.* (2006); 5, Turner *et al.* (2005); 6, Yan *et al.* (2006); 7, Casas *et al.* (2011); 8, Becana *et al.* (2020).

**Figure 4.**
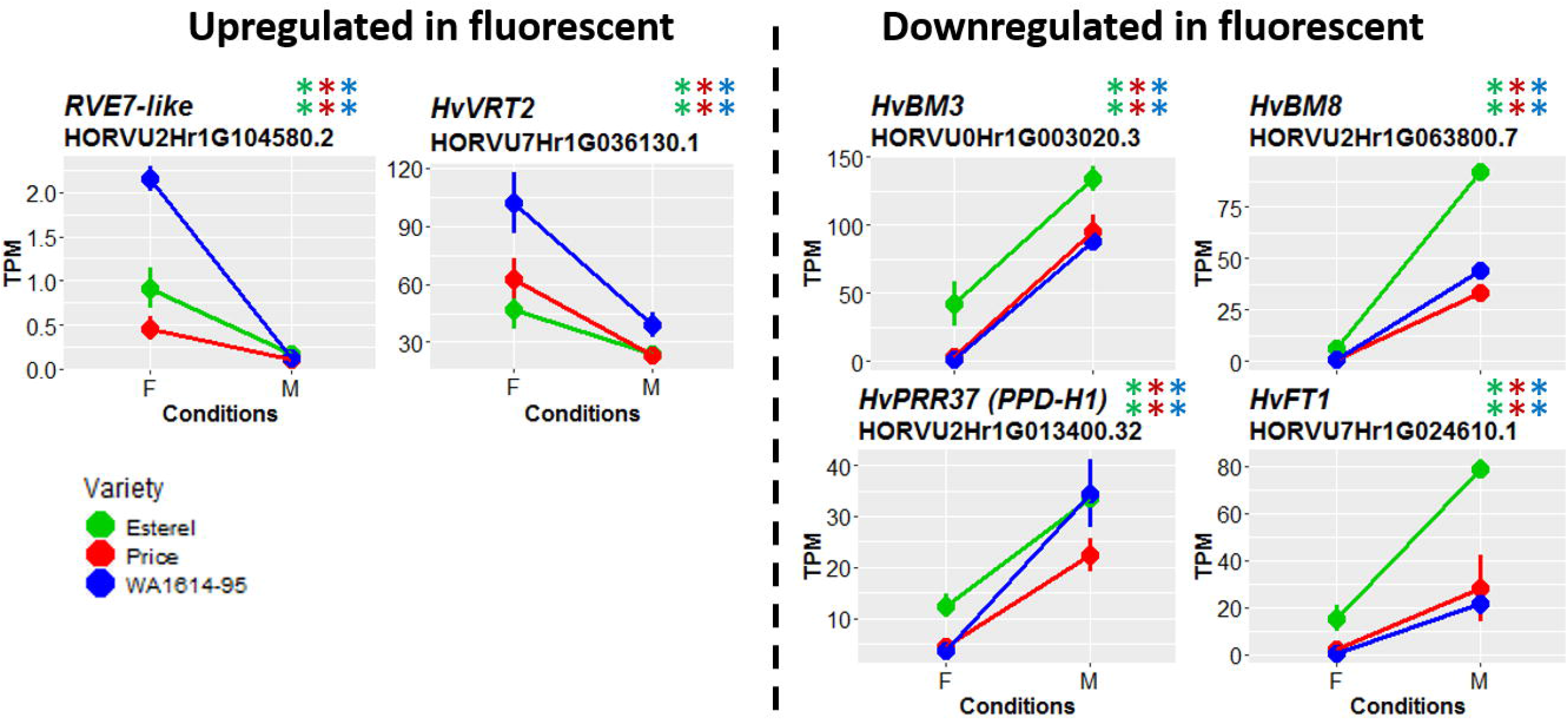
Expression levels of selected key genes belonging to flowering pathways, with similar expression patterns in the three genotypes. Two asterisks of the same colour indicate that differences are significant at q<0.01. Genes are separated in two groups: upregulated (left) and downregulated (right) under fluorescent conditions. Genotypes are coded as green (Esterel), red (Price) and blue (WA1614-95), in fluorescent (F) or metal halide conditions (M).

When we looked into key DE genes shared by just two varieties (Supplementary Dataset S1), Price and WA1614-95 (both considered sensitive lines to light quality) had the largest number (125, 66 up-regulated and 59 down-regulated in F, all in the same direction for both genotypes). In the set of DE genes shared by Esterel and Price, 20 genes showed differential expression with the same sign (14 up and 6 down), and 3 genes were different (up-regulated in Price and down-regulated in Esterel). Among the DE gene set shared by Esterel and WA1614-95, there were 13 genes with similar, and 29 with opposite trends. WA1614-95 had more up-(35) than down-regulated (7) genes, whereas Esterel had more down-(32) than up-regulated (10).

In the intersection between Price and WA1614-95, several transcripts coding for jasmonate-induced proteins were commonly up-regulated in condition F. When focusing on the DE genes in the intersection between WA1614-95 and Esterel, two WRKY family TFs with q value<0.01 were identified (Supplementary Dataset S1). With the initial q-value threshold of 0.05, naturally more genes were found (Supplementary Dataset S2), mostly up-regulated in F in the sensitive genotype WA1614-95 and down-regulated in the insensitive Esterel. Also, MADS-box TFs appeared frequently in the list of DE genes (as mentioned previously, *HvVRN1, HvVRT2, HvBM3, HvBM8, HvBM10, HvOS2*). Among genes annotated as MADS-box TFs, *HORVU3Hr1G095090* displayed the most extreme contrasting pattern of expression between sensitive (up-regulated) and insensitive (unchanged) varieties. BLASTN searches against NCBI nt and Ensembl Plants (Howe *et al*., 2020) databases revealed high similarity of this gene with a member of the wheat *FLC* subclade (*TaFLC-A4-2*, Schilling *et al*., 2020), and with *HvOS2* (*HORVU3Hr1G095240*), a neighbor gene on chromosome 3H. The latter has similar expression patterns (see section *Relevant light-response and developmental genes*), which might indicate a tandem duplication of *FLC*-like genes.

### GO enrichment analysis

We carried out a GO enrichment analysis to find functional commonalities among the genes present in each of the three sets of within genotype DE genes through shared GO terms. Within genotype DE genes calculated were independently subjected to GO enrichment analysis with “*de novo* assembly” and “Tibetan transcripts” as references. For robustness, the intersection of the resulting terms was declared as the final set of enriched GOs for each within genotype comparison (Supplementary Dataset S6). The main functional terms associated with DE genes are represented in Figure 5, with those involved in responses relevant for this study summarized in Supplementary Figure S3. Overall, the GO terms of DE genes suggest that major changes among varieties are related to translation and diverse metabolic processes. Those terms were associated with genes up-regulated in WA1614-95 and down-regulated in Price and Esterel (Figure 5A). The generic term “translation” also belongs to many biological processes already represented among the most significant GOs (organic substance metabolism process, cellular protein metabolic, cellular macromolecule biosynthetic, amide biosynthesis process, etc.). Thus, translation seems to be the main process underlying the extreme sensitivity of WA1614-95. Genes annotated with this GO category (n=153), mostly up-regulated, were related to ribosomal proteins in WA1614-95 (Supplementary Dataset S4), contrasting with the 37 and 121 DE genes found in Price and Esterel, respectively, mostly down-regulated in fluorescent light (Supplementary Datasets S3 and S5). Furthermore, the responses to light spectra seem variety-specific. Down-regulated DE genes in F were specifically associated with photosynthesis and responses to light stimulus and radiation in WA1614-95, to chloroplast organization in Price, and to organic substances metabolism and vacuolar activity in Esterel (Figures 5A and 5B).

**Figure 5.**
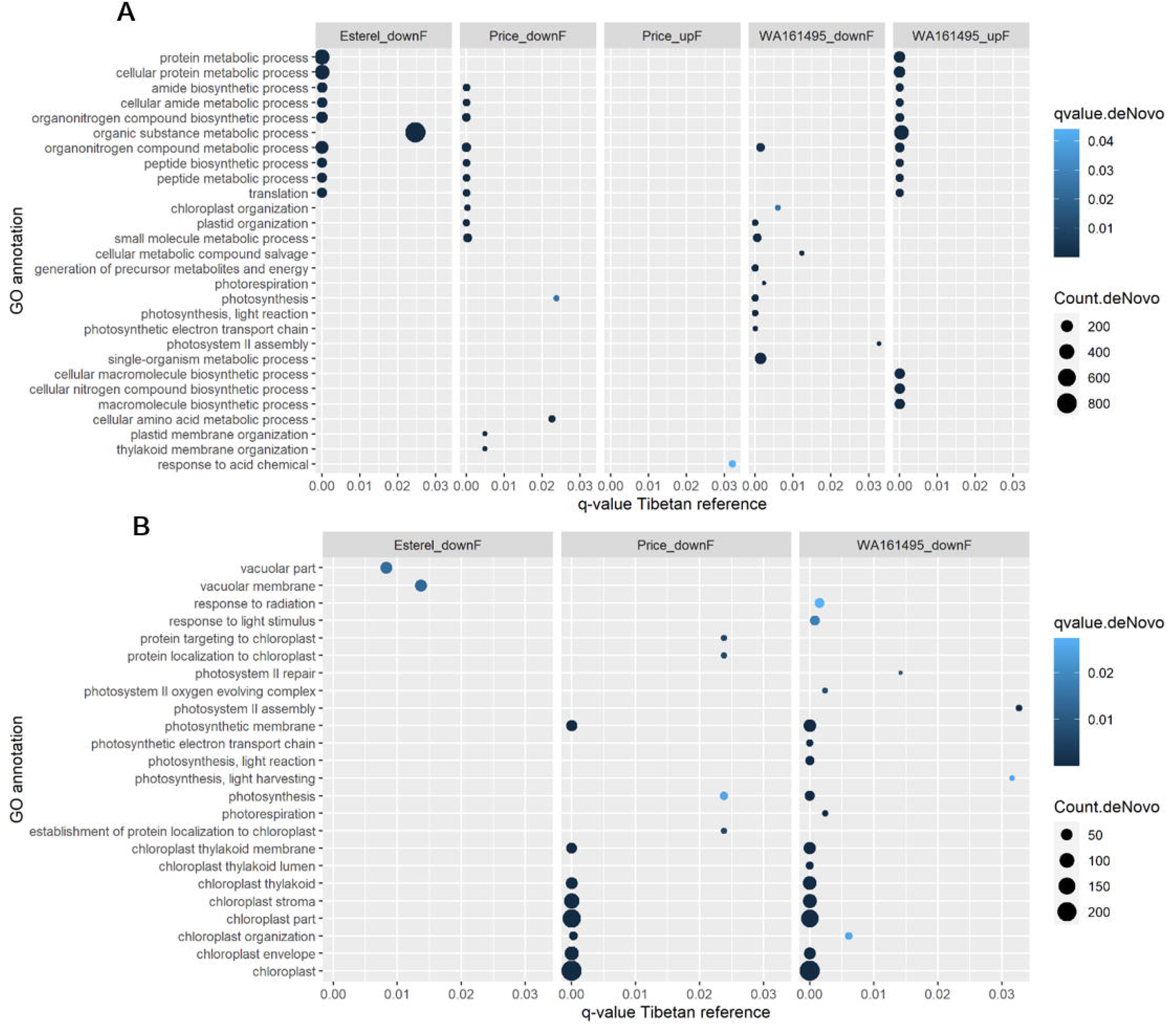
Enriched GO terms (q-value < 0.05), selected with different criteria. A) 10 most significant GO terms for each set of up- and down-regulated genes, ordered by increasing q-value in *de Novo* reference. B) Relevant GO terms for plant responses to light quality conditions. In both cases, only enriched GO terms corresponding to biological process (P) were selected. X-axis represents the q-value of the GO terms when taking the “Tibetan” reference, in Esterel, Price and WA1614-95 for each comparison: upF (upregulated in fluorescent conditions) downF, (downregulated in fluorescent conditions). Dot size is proportional to the number of transcript counts annotated to that GO term in the correspondent DE list.

We then relaxed the selection criteria of GO enrichment (p-value < 0.05), to have a wider look at categories involving developmental and light-specific responses (Supplementary Figure S3). Photosynthesis and responses to radiation, and abiotic stimulus, among others, were up-regulated in F in the insensitive genotype and down-regulated in the sensitive genotypes. Responses to starvation (down-regulated in F) and reproductive development (up-regulated in F) were only enriched in Esterel, whereas the response to red or far-red light appeared solely up-regulated in F in Price, and responses to UV-A and oxidation, and photosynthesis GOs appeared exclusively in WA1614-95 (down-regulated in F).

### Clusters of DE genes

To determine genes with possible common regulation, the Morex CDS DE genes (q-value<0.05), were grouped based on their expression patterns (see Figure 2). We created three sets of clusters, one for each of the three ‘within genotype’ DE gene sets. The optimal number of clusters was 39 for Price, 30 for Esterel, and 37 for WA1614-95. The clusters revealed different patterns of expression (Figure 6). Among them, some grouped genes with marked downregulation in F exclusively in the most sensitive variety WA1614-95 (Figure 6A, 6B). Other clusters grouped genes up-regulated in F in the sensitive varieties while showing stable expression in the insensitive (Figure 6C and 6D).

**Figure 6.**
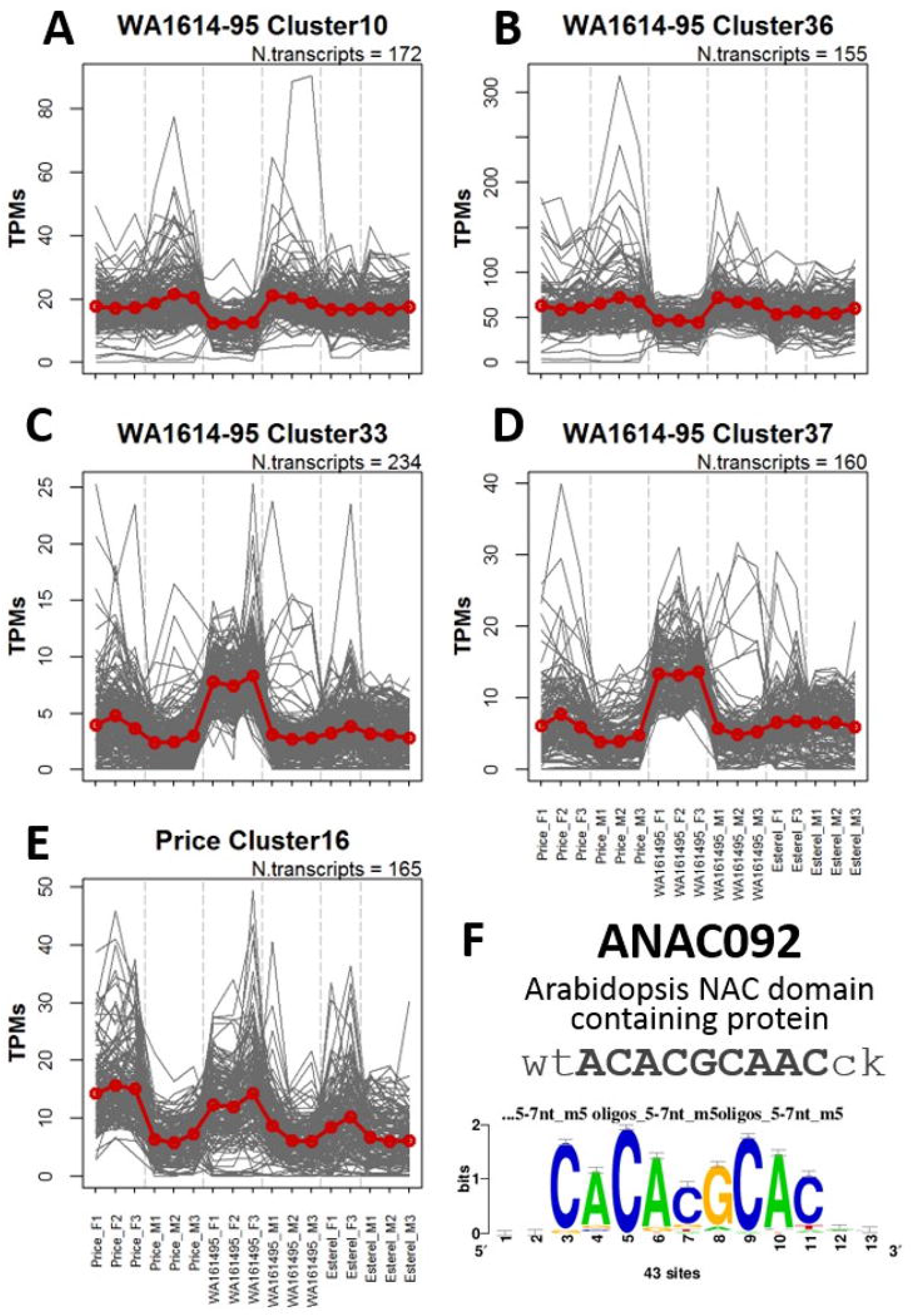
Relevant clusters of DE genes sharing expression patterns under fluorescent (F) and metal halide (M) conditions. Each grey line corresponds to the expression value of one gene across the different lines and conditions. Red lines indicate the average expression value of the sequences. (A, B) Clusters showing expression differences across light treatments in the most sensitive variety in F (WA1614-95). (C, D) Clusters showing expression differences across light conditions in sensitive (WA1614-95 and Price) varieties. (E) Cluster of genes showing a concurrent pattern of expression across light conditions in the three varieties. The number of genes within each cluster is indicated above each plot. (F) Common motif domain overrepresented in genes belonging to clusters 33 and 16. The consensus sequence was discovered in an upstream region covering [-500, 200] bp. The significance of the motif is proportional to the height of the letters. The consensus sequences found are similar to that of *Arabidopsis thaliana* ANAC092 annotated in footprintDB.

Cluster 16 from the Price DE genes showed a common up-regulation in F in the three varieties (Figure 6E). The 5 clusters highlighted (Clusters 10, 33, 36, 37 in WA1614-95 and Cluster 16 in Price) were subjected to a motif discovery analysis. Upstream sequences of genes within clusters 16 and 33 showed enriched DNA conserved motifs similar to *Arabidopsis* ANAC092 (Figure 6F), suggesting that a barley homolog of ANAC092 could be coordinating the expression of the genes within these clusters.

### Relevant light-response and developmental genes

A large number of DE genes were found (Supplementary Datasets S3, S4, and S5) beyond those in Table 1. We narrowed down the list focusing on genes known to be involved in light perception (phytochromes, cryptochromes), circadian clock, flowering initiation, and development (Figure 7). Among these, we found that *HvPhyC* was up-regulated in F in the three genotypes, whereas *HvPhyB* and *HvCry2* were only differentially expressed in Price (up-regulated in M), whilst no differences were found for *HvPhyA* or *HvCry1a* (Supplementary Figure S2). Two TFs that act downstream in the photoreception machinery and the light-signal transduction, *HvPIF5*, and *HvHY5*, were expressed with the same pattern as *HvPhyC*.

**Figure 7.**
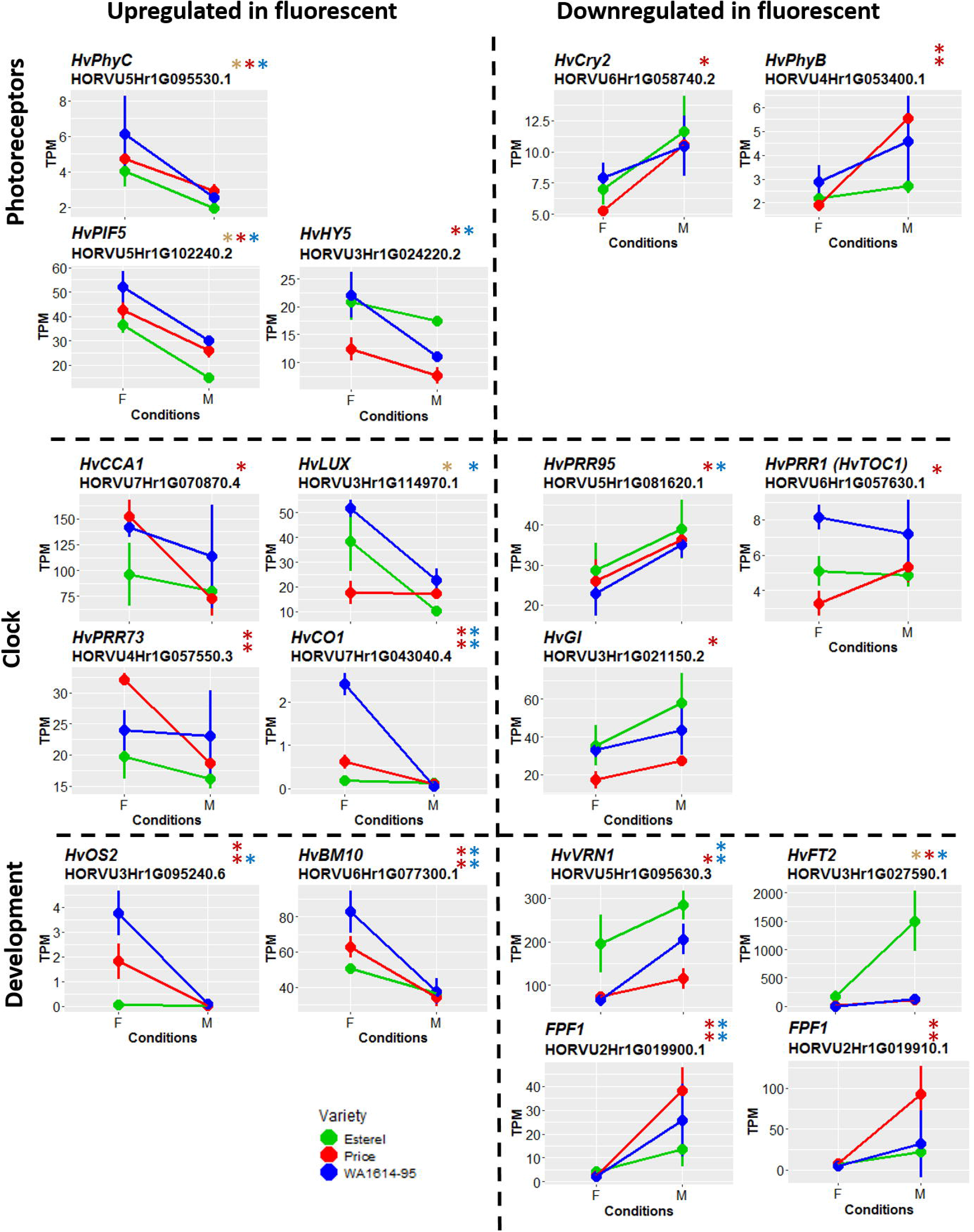
Expression levels of flowering-related genes (photoreceptors, circadian clock and development) in the three genotypes. Genes are separated in two groups: upregulated (left) and downregulated (right) under fluorescent conditions. F, fluorescent conditions; M, metal halide conditions. Genotypes coded as in Figure 4. Differences between treatments are significant at q-value < 0.05 (^*^) or q-value<0.01 (^**^).

The three genotypes showed reduced transcription levels of *PPD-H1, HvFT1* (Figure 4), and *HvVRN1* (Figure 7) in fluorescent conditions, consistent with the delayed plant development. Esterel showed higher expression levels than the sensitive genotypes for *HvFT1* and *HvVRN1*, in accordance with its accelerated development in both conditions. Besides, the three genotypes showed increased transcript levels of the flowering repressors *HvOS2* (Figure 7), *HvVRT2*, and an orthologue of *RVE7*-like in wheat under fluorescent light (Figure 4). WA1614-95, which was the latest flowering genotype, showed the highest transcript levels of these repressor genes. On the other hand, *HvFPF1*-like (*Flowering Promoting Factor 1*) transcripts were up-regulated in Price and WA1614-95 under M.

We also identified some differentially expressed genes, mainly in Price and some of them in WA1614-95, that encode components of the circadian clock: *HvCCA1, HvLUX*, and *HvPRR73* up-regulated in F; *HvGI, HvTOC1*, and *HvPRR95*, down-regulated in F; and clock output genes, as *HvCO1*, up-regulated in F.

Two members of the *C-REPEAT/DREB BINDING FACTOR* (CBF) family (*HvCBF14, HvCBF4a*), one member of the *COLD-RESPONSIVE* (COR) family (*WCOR15A*) and one from the *INDUCER of CBF EXPRESSION* (*ICE*), an ortholog of rice (*ICE-like* annotated as *metacaspase I*), all relevant in the acquisition of freezing tolerance, were up-regulated in fluorescent light, mainly in the sensitive genotypes (Figure 8).

**Figure 8.**
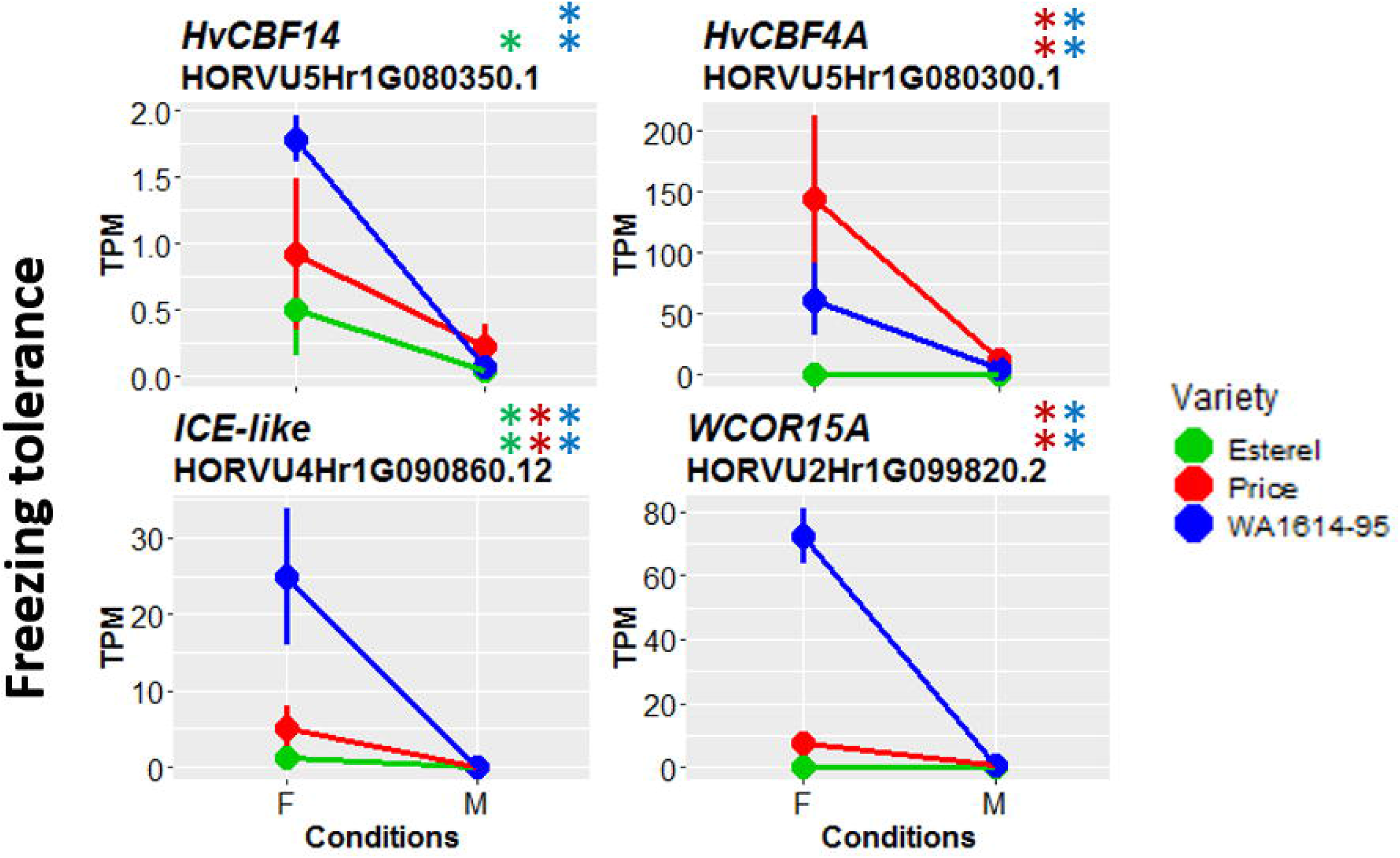
Expression levels of genes related to frost response in the three genotypes. Genes are separated in two groups: upregulated (left) and downregulated (right) under fluorescent conditions. F, fluorescent conditions, M, metal halide conditions. Genotypes coded as in Figure 4. Differences between treatments are significant at q-value < 0.05 (^*^) or q-value <0.01 (^**^).

The RNA-seq results were validated through qRT-PCR analysis using 10 genes responsive and non-responsive to light quality conditions (Figure 9). Samples were extracted from the same genotypes, conditions, and age in an independent experiment. We obtained a positive Pearson correlation (r = 0.70).

**Figure 9.**
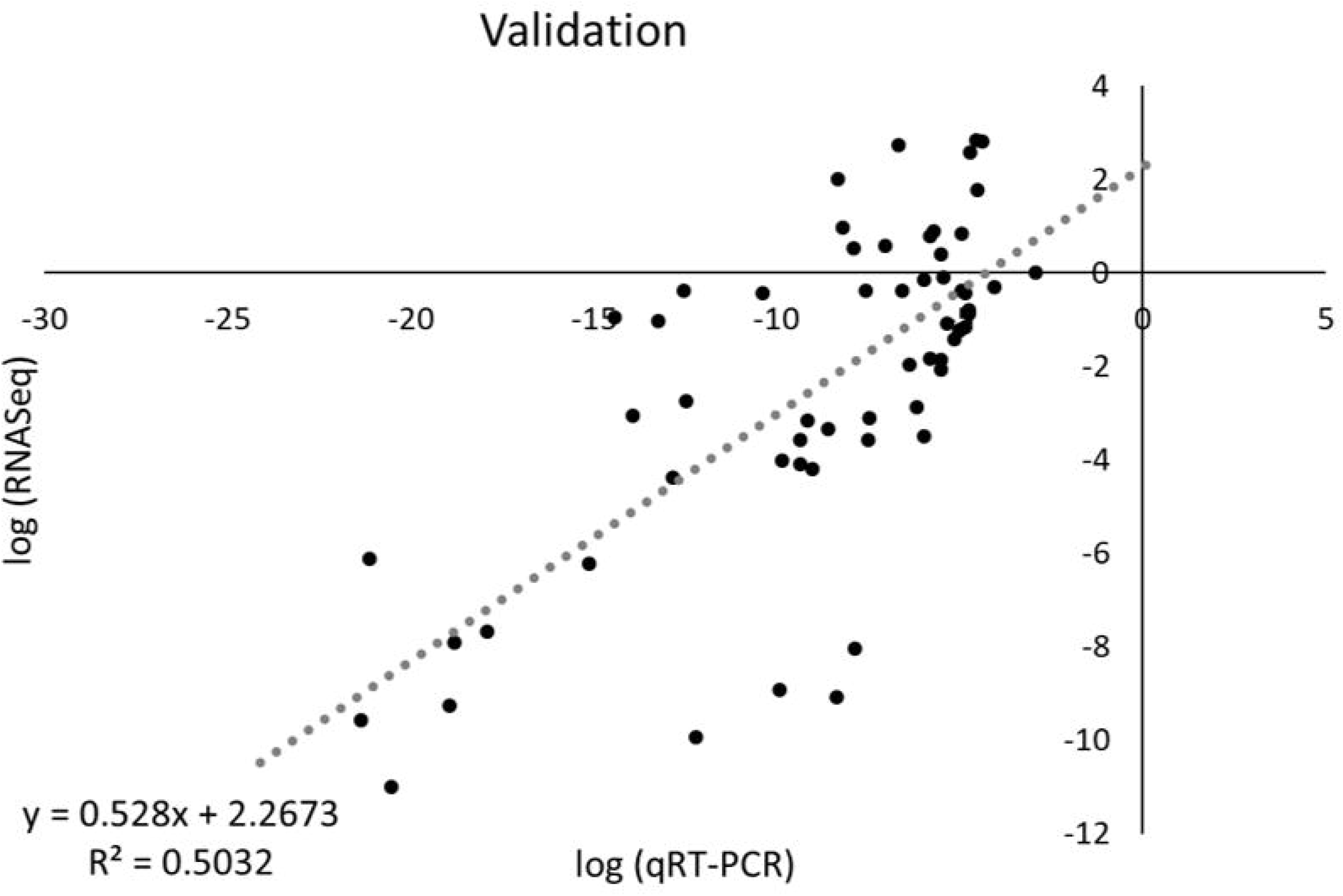
Correlation between RNA-seq and qRT-PCR expression values. Values obtained after normalization (Log2 of gene expression relative to *Actin*). The dotted line corresponds to a linear regression. R^2^, coefficient of determination.

## Discussion

### Light quality affects to development genes

Delayed development of fluorescent light-grown plants was paralleled by dramatic changes in the expression of development-related genes, which were overrepresented among the 14+1 key DE genes. Among them, the flowering repressors *RVE7-like, VRT2*, and *ICE-like* were up-regulated in fluorescent light, whereas *AP1* MADS-box photoperiod responsive genes *HvBM3* and *HvBM8*, and *HvFT1* were down-regulated. The large reduction of expression of *PPD-H1* could explain the downregulation on these last three genes because it mediates the long-day induction of *HvFT1* (Turner *et al*., 2005), and several studies have reported the *PPDH1*-dependent up-regulation of *HvBM3* and, *HvBM8* during development (Digel *et al*., 2015, 2016; Ejaz and von Korff, 2017).

GO analysis of DE genes did not suggest the existence of general responses to light spectra. Differences in overrepresented GO terms among the three varieties indicated the presence of different light quality responses. For instance, translation and diverse metabolic processes were associated with up-regulated genes in the sensitive-extreme variety and down-regulated in the other two (Figure 5A). Ribosomal protein genes were overrepresented in the sensitive-extreme variety, indicating a transcriptional reprogramming or translational regulation only in this genotype, as found in other species subjected to biotic and abiotic stresses (Solano-De la Cruz *et al*., 2019). Furthermore, genes down-regulated in F were associated with chloroplast and plastid organization and photosynthesis responses in the sensitive varieties only, and not in Esterel (Figure 5; Supplementary Figure S3). Therefore, the fluorescent light spectrum might alter the photosynthetic electron transport chain differently in these barley varieties. In conclusion, the different grades of sensitivity to light spectra were associated with transcriptional reprogramming, plastid signals, and photosynthesis. Interestingly, plant development reprogramming in response to high light intensity in *Arabidopsis thaliana* has been related to epigenetic changes involving *FLC* activity (Feng *et al*., 2016). Fittingly, *HvOS2*, the barley *FLC* orthologous gene, was up-regulated in F in both sensitive varieties.

### Sensitive varieties experienced partial reversion of vernalization and displayed cold tolerance responses

Sensitive and insensitive varieties showed strikingly different patterns of DE genes. However, there was a remarkable similarity in the sets of DE genes for sensitive varieties WA1614-95 and Price.

TFs seemed overrepresented among the DE genes. Although some MADS-box TFs were equally affected across varieties (*HvBM3, HvBM8, HvVRT2*), other MADS-box genes showed contrasting patterns following the varieties’ sensitivity. This was the case of *FLC*-like *HvOS2* and its paralog *HORVU3Hr1G095090*. In wheat, the duplication within the *FLC*-clade has been related to adaptation (Schilling *et al*., 2020). *HvOS2* represses the expression of *Flowering Promoting Factor1*-like genes (Greenup et al. 2010; Hemming et al. 2012), which also appear differentially expressed in our study (Figure 7). *HvOS2* expression responds to cold, mediated by *HvVRN1*, which was also differentially expressed only in the sensitive varieties. This gene should have been fully induced after vernalization in all three varieties (even more so in WA1614-95, which needs little vernalization), but it was less induced in F light in Price and WA1614-95, not different from a light-mediated de-vernalization.

The high expression of *HvOS2* and its nearby paralog, as well as other genes related to cold acclimation (*HvCBF14* and *WCOR15a*) under F light in sensitive varieties, bodes well with their reduced *HvVRN1* expression because all these genes are VRN1 targets in barley (Deng *et al*., 2015). Cold-acclimation responses elicited by light are not a new finding. Novák *et al*. (2016) reported that barley plants grown under fluorescent light supplemented with far-red light presented high *HvCBF14* induction, increasing their freezing tolerance, but these results were found in plants during the hardening process, not in fully vernalized plants grown under inductive conditions, as was the case here. We also found two members of the CBF-clade (*HvCBF14* and *HvCBF4a*, Skinner *et al*., 2005) up-regulated in fluorescent conditions. A relationship between regulation of the *CBF* regulon and light (low R:FR ratio), mediated through phytochromes, and under higher temperatures than those that confer cold acclimation, was reported by Franklin and Whitelam (2007). In a related manner, freezing tolerance genes were affected in *phyB*-null mutants in wheat (Pearce *et al*., 2016), rice (He *et al*., 2016), and *Arabidopsis* (Franklin and Whitelam, 2007), causing downregulation of a member of the *INDUCER of CBF EXPRESSION* (*ICE*) gene family (Badawi *et al*., 2008). All these pieces of evidence strongly suggest a role of *PhyB* in light-mediated activation of cold acclimation pathway, and our results support this hypothesis.

Adding to the cold-like effect of the fluorescent light, genes related with cold acclimation as *VRT2* (Kane *et al*., 2005), a homolog of *RVE-7*, and an *ICE*-like protease showed consistent higher expression under fluorescent light. We hypothesize that upregulation of repressors and cold-induced genes under fluorescent light in fully-vernalized plants and LD indicate that these plants are not sensing the favorable conditions, and remain in the cold acclimation phase, eliciting cold-related responses, particularly in the sensitive varieties.

### Sensitive varieties experienced phenomena related to senescence

Among the highly significant DE genes, the only one showing opposite directions between insensitive Esterel (down) and the sensitive varieties (up) was an *ethylene-responsive transcription factor*. Genes encoding for jasmonate-induced proteins were up-regulated in fluorescent light in the sensitive varieties. Jasmonate and ethylene are hormones involved in plants’ responses to a wide range of abiotic stresses, and the latter is also involved in senescence. This might be connected with *hemoglobin 1*, a key gene up-regulated under fluorescent light in the three varieties. Non-symbiotic hemoglobins are involved in abiotic stress responses, with a purported role in protecting cells from dehydration by modulating nitric oxide concentration (Rubio *et al*., 2019; Becana *et al*., 2020). Sensitive varieties were suffering from abiotic stress under fluorescent light, and these pathways are good candidates to explain the phenotypic reactions of sensitive and insensitive varieties.

The clusters of DE genes up-regulated in fluorescent light in sensitive varieties were enriched in regulatory motifs similar to *ANAC092*, a regulator of senescence, belonging to the *NAC* family (Balazadeh *et al*., 2010). NAC TFs control multiple processes, although they are mainly associated with senescence and response to abiotic stresses, integrating also cold signals and flowering (Yoo *et al*., 2007). The set of DE genes shared by WA1614-95 and Esterel showed mostly opposite trends of expression, in accordance with their sensitivities. Among them, there were several encoding for WRKY TF. These TFs are involved in the regulation of transcriptional reprogramming associated with plant abiotic and biotic stress responses (Bakshi and Oelmüller, 2014) and, at least in grapevine, WRKY TFs trigger cell wall modifications to block the entrance of UV light into the cell (Lesniewska *et al*., 2004).

Interestingly, a link between cold acclimation, senescence, and flowering delay was described by Wingler (2011) for *Arabidopsis thaliana* and barley. Through a promoter motif analysis, this author found that flowering and senescence regulation are closely associated with light and stress signaling and that cold-responsive genes were induced in plants with delayed senescence.

### Signaling pathways affected by light quality

The variability of expression patterns found for genes related to response to light and specific spectra regions (red, far-red, UV) indicates that the treatments affected the function of light receptors and photomorphogenesis. We found higher expression levels of *HvPhyC* under fluorescent light in all three genotypes, and an opposite trend for *HvPhyB* and *HvCry2* (consistent with their antagonist role, reported by Más *et al*., 2000). According to our results, the fluorescent light caused a strong imbalance of the expression of these three signaling genes, which could lead to a disruption of 1) the balance of active and inactive forms of phytochromes, 2) patterns of occurrence of homo- and heterodimers, and 3) phytochromes to cryptochromes ratios. These imbalances could be at the top of the cascade of changes in gene expression found in downstream pathways.

Both PhyC and PhyB are required for the photoperiodic induction of flowering in wheat (Pearce *et al*., 2016; Kippes *et al*., 2020). Null wheat mutants of either *PhyC* or *PhyB* produced late-flowering plants, with altered vegetative development. Up-regulation of *HvPhyC* in F light resembled the phenotype of lacking a functional gene in wheat.

PhyC signal may have cascaded down through PPD-H1, as it is known to activate PPD-1 in LD (Nishida *et al*., 2013; Chen *et al*., 2014; Woods *et al*., 2014). In our study, there were opposite patterns of expression between the DE genes *HvPhyC* and *PPD-H1* in the three genotypes, supporting their close relation. The altered phytochrome expressions may have shifted their proportions from the optimum, reducing *PPD-H1* induction in fluorescent light.

Also, several signaling and clock genes were DE in the sensitive lines. In *A. thaliana*, Franklin and Whitelam (2007) observed that the phytochrome signaling was mediated by the circadian clock. Thus, the complex relation between photoreceptors and the clock might provide a reason for the differences in downstream development genes, and in the phenotype. It is remarkable that *HvPhyB, HvCry2*, and most of the clock genes represented, were only differentially expressed in cultivar Price.

### Why would an annual crop develop mechanisms of adaption to light quality?

We have revealed the presence of phenotypic variation of barley in response to light quality. Could this variation be adaptive? Barley spread from its cradle at 35°-40°N to latitudes beyond 60°N involving well-known adaptation mechanisms like insensitivity to day length (Jones *et al*., 2008). Adaptations to other possible light-related factors for crop species have been largely overlooked. If factors other than day length underlie light adaptation of crops, this area deserves urgent attention. Climate change is already causing latitudinal shifts of variety distribution and plant breeders should know how to cope with possible light-related genetic effects other than photoperiodic response (Hunt *et al*., 2019).

Light quality effects have received more attention in perennials. In a recent report, Chiang *et al*. (2019) found that light quality affects tree growth showing a wide natural variation. This is caused by the latitudinal change in the duration of periods under low solar angles and by variation of overcast conditions (that implies variable R:FR ratios). They suggested that trees that originated at high latitudes are more sensitive to light quality, compared to those from lower latitudes. The barley lines tested in our experiment are autumn sown, and much of their growth period occurs in winter, the period of the lowest solar angle. It is conceivable that annual crops also took advantage of mutations to optimize their growth in regions with light quality features different from the ones found at their center of origin. In addition, the solar angle in early spring is still relatively low and shows a large latitudinal variation. This may have resulted in the evolvement of additional regulation mechanisms of plant development at the higher latitudes, preventing the precocious initiation of stem elongation in fully vernalized plants.

To conclude, we have found considerable variability among barley genotypes regarding light quality sensitivity, involving different molecular mechanisms, akin to other abiotic stress responses. Further research is needed to pin down their molecular bases and, in particular, the extent of their role under natural conditions, with possible repercussions on crop breeding.

## Abbreviations

DE: differentially expressed
DEV: developmental stage
F: fluorescent
LD: long day
M: metal halide
TF: transcription factor

## Supplementary data

### Supplementary Tables and Figures

**Supplementary Table S1**. List of the barley genotypes examined and allelic variants for the major genes of flowering time.

**Supplementary Table S2**. Primer sequences for qRT-PCR assay.

**Supplementary Figure S1**. Correlation of transcript abundances across biological replicates.

**Supplementary Figure S2**. Selection of flowering related genes (photoreceptors, circadian clock and development).

**Supplementary Figure S3**. Enriched GO terms relevant for plant responses to light quality conditions.

### Supplementary Information

**Supplementary Information S1**. Summary of the DE experimental design and results of DE genes from within line comparison.

**Supplementary Information S2**. Number of differentially expressed genes (DE) in each of the six between line comparisons.

**Supplementary Information S3**. Number of biological samples correctly clustered in the hierarchical clustering generated for the nine comparisons.

### Supplementary Dataset

**Supplementary Dataset S1**. List of Key DE genes (q-value< 0.01) shared between varieties.

**Supplementary Dataset S2**. List of DE genes (q-value < 0.05) shared between varieties.

**Supplementary Dataset S3**. List of DE genes in Price, comparison between M and F conditions.

**Supplementary Dataset S4**. List of DE genes in WA1614-95, comparison between M and F conditions.

**Supplementary Dataset S5**. List of DE genes in Esterel, comparison between M and F conditions.

**Supplementary Dataset S6**. GO enrichment.

## Acknowledgments

This work was supported by the Spanish Ministry of Economy and Competitiveness (Projects AGL2013–48756-R, including a scholarship granted to AM, and AGL2016– 80967-R), and Spanish Ministry of Science and Innovation (PID2019-111621RB-I00). The authors acknowledge Najla Ksouri and Francesc Montardit for their technical support.

## Author contributions

AC, BC-M, EI, IK: Conception and design.

BC-M, AR-R: Methodology

AR-R, AM: Formal analysis, Writing - Original draft.

AM: Visualization

TK, MM, IK: Acquisition of data

AC, EI: Writing – Review & Editing, Project administration, Funding acquisition

AC, EI, BC-M: Supervision

## Data availability

The data for this study have been deposited in the European Nucleotide Archive (ENA) at EMBL-EBI under accession number PRJEB35759 (https://www.ebi.ac.uk/ena/browser/view/PRJEB35759). Additionally, motif discovery analysis of selected clusters of differentially expressed genes is available at rsat.eead.csic.es/plants/data/light_report/.

